# Fat SIRAH: Coarse-grained phospholipids to explore membrane-protein dynamics

**DOI:** 10.1101/627570

**Authors:** Exequiel E. Barrera, Matías R. Machado, Sergio Pantano

## Abstract

Tne capability to handle highly heterogeneous molecular assemblies in a consistent manner is among the greatest challenges faced when deriving simulation parameters. This is particularly the case for coarse-grained simulations in which chemical functional groups are lumped into effective interaction centers for which transferability between different chemical environments is not guaranteed. Here we introduce the parameterization of a set of CG phospholipids compatible with the latest version of the SIRAH force field for proteins. The newly introduced lipid species include different acylic chain lengths, partial unsaturation, as well as polar and acidic head groups that show a very good reproduction of structural membrane determinants, as areas per lipid, thickness, order parameter, etc., and their dependence with temperature. Simulation of membrane proteins showed unprecedented accuracy in the unbiased description of the thickness-dependent membrane-protein orientation in systems where this information is experimentally available (namely, the SarcoEndoplasmic Reticulum Calcium –SERCA-pump and its regulator Phospholamban). The interactions that lead to this faithful reproduction can be traced down to single amino acid-lipid interaction level and show full agreement with biochemical data present in the literature. Finally, the present parameterization is implemented in the GROMACS and AMBER simulation packages facilitating its use to a wide portion of the Biocomputing community.

## INTRODUCTION

A typical eukaryotic cell membrane contains cholesterol and several hundreds of different lipid species including phosphatiylcholines (PC), sphingomyelin, and gangliosides, which are more abundant in the the outer leaflet; and phosphatidylethanolamines (PE) and anionic species like phosphatidylserine (PS), phosphatidic acid and phosphatidylinositol typically found in the inner leaflets.^1^ The lipid content of different biological membranes modulates not only their own biophysical characteristics but also their interactions with proteins. Such interactions determine the functional orientation of proteins and their sorting into different organelles.^2^ Big efforts in modeling these heterogeneous systems have been done in the last years, being worthy to mention the work performed by Ingólfsson et al. in 2014 to model a mammalian plasma membrane composed by nearly 20000 lipids of 63 different types.^3^ Concurrently, the increasing knowledge about bacterial lipid composition has encouraged the modeling community to generate parameters for this particular membranes, such is the case of the atomistic force field CHARMM which contains not only the most common lipids found in higher organisms but also an extensive set of lipopolysaccharide fragments.^4^

Currently, the Orientations of Proteins in Membrane database (OPM) (https://opm.phar.umich.edu) contains more than 4 thousand structures constituting a prized source of information for protein-lipid interactions applicable in the generation of optimized parameters.

Despite the success of theoretical techniques to reproduce the structure and dynamics of membrane related systems, their size and complexity still impose enormous computational costs. This has prompted the development of coarse-grained (CG) models that dodge this issue by reducing degrees of freedom not resigning chemical specificity. Starting with the pioneering work by the group of Michael Klein,^5^ several groups have studied in a simplified fashion the structure and dynamics of lipid species using implicit^6,7^ or explicit solvent models^8^. Nowadays the most complete repertoire is provided by the MARTINI force field, which features over 180 different species^9^ and has been recently implemented to work both in explicit and implicit solvent.^10^ Less common, however, is the availability of CG parameters to work in combination with proteins or other biological entities. For state of the art in biological membrane modeling and lipid-protein interactions, see the reviews by Marrink et al.^11^ and Corradi et al.^12^

Here, we present a general parameterization for phospholipid molecules at CG level, developed to work with the latest version of the SIRAH force field for proteins and explicit solvent.^13^ The set of lipids represented includes dimyristoyl phosphatidyl choline (DMPC), dipalmitoyl phosphatidyl choline (DPPC), palmitoyl oleoyl phosphatidyl choline (POPC), palmitoyl oleoyl phosphatidyl ethanolamine (POPE) and palmitoyl oleoyl phosphatidyl serine (POPS). This diversity is further enriched by the additional implementation of fragment-based topologies allowing for the easy combination of PC, PE and PS heads with myristoyl (MY), palmitoyl (PA) and oleoyl (OL) tails to create new lipids. Our phospholipid selection provides a variety of polar and charged head groups, lengths and unsaturation degrees of acyl chains, which minimally covers representative membrane compositions found in mammalian cells and biophysical assays.^14,15^

Following a top-down approach, parameters were validated against a series of structural and dynamic properties experimentally measured on planar bilayers. These include area per lipid, thickness, average acyl chain order, lateral diffusion coefficient and density profiles along the perpendicular axis to the membrane surface. A series of simulations spanning a temperature range between 303-333 K, which covers the largest portion of biological applications, showed a very good reproduction of the experimental data available in the literature.

Finally, we assessed the compatibility with our recently updated force field for proteins (SIRAH 2.0)^13^ by addressing the dynamics of challenging membrane proteins as the staphylococcal Alpha-hemolysin (αHL) heptameric pore, the bacterial outer membrane protein OmpX, the Sarco Endoplasmic Reticulum Calcium ATPase (SERCA) and its regulator Phospholamban (PLB). We found a remarkable agreement with fundamental features reported for these proteins, as lipid dependent protein tilting, amino acid specificity for the so called “floating”, “snorkeling” and “anchoring”. Worth mentioning, this happens without resigning the capacity of SIRAH to work within the standard Hamiltonian used for MD simulations and reproduce secondary structures in absence of constraints or external biases. Additionally, the addresed test cases help to point out certain limitations of the force field that users should have in mind when choosing CG approaches to simulate their systems of interest. The present implementation for natively running MD CG simulations using GROMACS or AMBER engines is freely available from our web site (www.sirahff.com).

## METHODS

### Mapping to SIRAH force field

The SIRAH pipeline to run a MD simulation starts on mapping the atomistic coordinates of a system to its corresponding CG representation. This step is facilitated by SIRAH Tools^16^ through the use of so called MAP files, which provide the transformation rules for mapping the positions of atoms to CG beads. It may be worth recalling that beads in SIRAH use the same position of real atoms. The topology, which includes connectivity and parameters, is then generated from the CG coordinates by native ports implemented in GROMACS and AMBER packages. Transferability and usability was always a major concern in the developing of SIRAH. In that sense, lipids present an important challenge, as residue and atom names are not as commonly standardized as proteins or nucleic acids.^17^ Indeed, different conventions of three to four letter codes were adopted during the last years by force field developers, some of which being rather ambiguous, custom made or case specific.^18^ Usually, residue-based topologies are used to describe individual lipid molecules such as DMPC, DPPC, POPC and so on. However, starting from Lipid11 force field, a modular framework for lipid residues was implemented in the AMBER package.^19^ The use of fragment-based topologies to separately describe head and tail groups expanded the accessible molecular space with few parameters at the cost of resigning an easy lipid identification.

In the present work, the SIRAH package is extended to provide support for both residue- and fragment-based topologies for DMPC, DPPC, POPC, POPE and POPS molecules, PC, PE and PS head groups, and MY, PA and OL tails. MAP files are compatible with CHARMM 27/36,^20,21^ Berger lipids,^22^ GROMOS flavors (43a1,^23^ 43a1-s3,^24^ 53a6,^25^ 87^26^ and CKP^27^), OPLS-AA,^28^ OPLS-UA,^29^ GAFF,^30^ Slipids,^31^ Lipid11-17^32^ as they are implemented in Charmm-GUI (http://www.charmmgui.org),^33^ LipidBook (https://lipidbook.bioch.ox.ac.uk),^34^ Tieleman’s database (http://people.ucalgary.ca/~tieleman/download.html), GROMACS repository (http://www.gromacs.org) and AMBER libraries (http://ambermd.org). Despite this effort, users are advised to check and modify the MAP files as required.

Packing lipids and embedding proteins into membranes is other important issue. In addition to grant compatibility with outputs from Charmm-GUI, MemBuilder (http://bioinf.modares.ac.ir/software/mb), membrane builder extension of VMD (https://www.ks.uiuc.edu/Research/vmd/plugins/membrane), HTMD database (http://www.playmolecule.org/OPM),^36^ PDB files for building systems with Packmol^37^ are included. The updated SIRAH package for AMBER and GROMACS, including step-by-step tutorials, is freely available at http://www.sirahff.com.

### Derivation of the model

SIRAH uses a standard Hamiltonian common to most MD simulation packages, including bonds, angles, torsionals, electrostatics and van der Waals (vdW) terms. To ensure the compatibility of the newly created lipid representation with all the molecules already present in our force field we employed bead types recently introduced in the updated version of SIRAH (Figure 1).^13^ According to the SIRAH’s philosophy of reproducing structural determinants, parameters were adjusted to fit primarily areas per lipid and membrane thickness.

**Figure 1.**
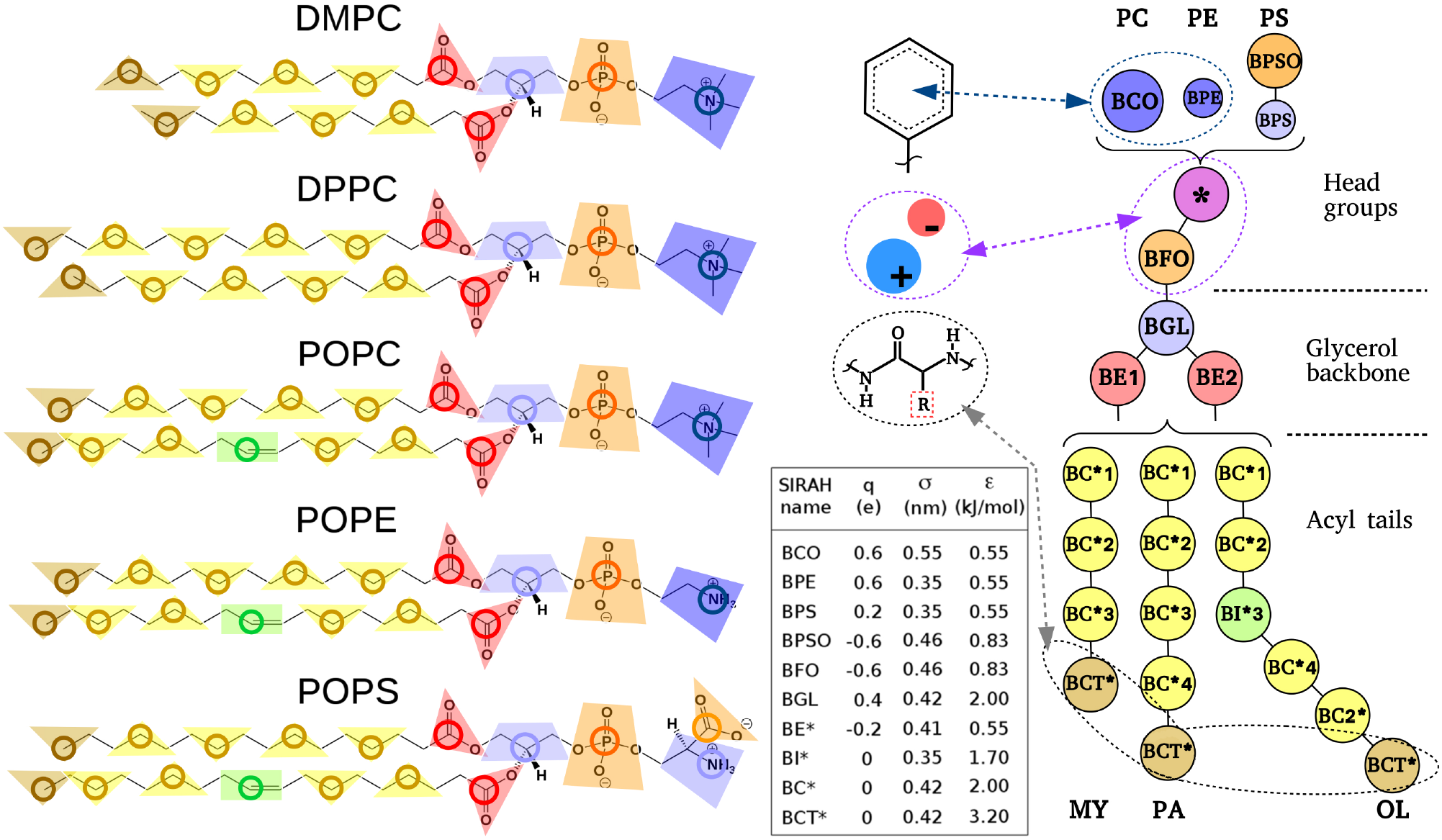
Schematic representation of CG mapping for SIRAH lipids. Non-bonded parameters for each bead are detailed in the table. Selected atomic positions used for mapping are highlighted with bold colored circles. All beads carry a uniform mass of 50 a.u., bonded with spring constants of 41,840 kJ mol^−1^ nm^−2^, and angle bending constants of 627.6 kJ mol^−1^ rad^−2^ and 14 kJ mol^−1^ rad^−2^ for polar heads and saturated acyl chains, respectively. Equilibrium distances and angles are taken from Shelley et al.^39^ vdW interactions between particular beads calculated outside the Lorentz-Berthelot combination rules (see text) are identified with dashed arrows and circles.

Topologies of the coarse grained DPPC, POPC, POPE and POPS are based on the previously reported DMPC model,^38^ Modeling of Palmitoyl acyl chains was simply done by adding one hydrophobic methylene-like bead to the myristoyl chains already existing for DMPC. Oleoyl chains required to model unsaturated cis-bonds setting the equilibrium angle value to 120° with an angle-bending constant of 20 kJ mol^−1^ rad^−2^. For this case, new terminal beads were created, keeping van der Waals values but shortening its equilibrium distance to 0.367 nm. This implementation should be considered in the hypothetical cases of creating new acyl chains containing carbon atoms multiple of three (e.g. Laureoyl). In addition to the existing bead representing a choline group, two new polar heads were created, namely ethanolamine- and serine-phosphatidyl groups. For ethanolamine (Figure 1) a positively charged bead (+0.6 e) with smaller size than the choline bead was chosen (σ = 0.35). For serine anionic heads, a bead with the same σ and ε values as ethanolamine but charged +0.2 e plus an extra negatively charged bead, accounting for the dipolar characteristics of the amine and carboxylate groups of this moiety. In this case, the sum of partial charges distributed in the beads of the phosphatidyl serine head equals to −1. Lorentz-Berthelot combination rules are used to calculate most of the vdW interactions. However, a reduced set of beads pairs use specific parameters outside those rules to enhance physicochemical specificity. In particular, three different types of vdW interactions between phospholipids and protein beads were calculated outside the Lorentz-Berthelot combination rules to ensure the correct lipid-protein interplay without perturbing membrane properties. First, interactions between polar heads (choline, ethanolamine and serine) and oppositely charged side chains beads were treated analogously to salt-bridges calculated for protein-protein interactions.^13^ Moreover, cation-π interactions are considered the same way as in proteins, and positively charged head groups utilize ε values of 1.32 kJ mol^−1^ when interacting with aromatic side chains. Lastly, the terminal beads (named BCT* in Figure 1) were assigned a vdW well corresponding to ε = 3.2 kJ mol^−1^ to ensure the acyl chain interdigitation, which impacts in the correct reproduction of structural properties (see *Pure lipid simulations* section in Results). However, the interaction between the terminal acyl beads and hydrophobic protein beads was set to ε = 2.00 kJ mol^−1^, mimicking the interaction of the methyl-like group used to parameterize Alanine.^13^ A complete description of non-bonded interaction parameters is depicted in Figure 1.

### Computational details

Initial bilayer configurations were built with Packmol.^37^ Simulations were carried out using GROMACS 2016.1^40^ (http://www.gromacs.org) and/or AMBER16 (http://ambermd.org) GPU codes^41,42^. A direct cut-off of 1.2 nm was chosen for non-bonded interactions and long-range electrostatics were calculated using the Particle-Mesh Ewald method.^43^ Solvation was done using a pre-equilibrated box of SIRAH’s CG water molecules (named WT4).^44^ After energy minimization, achieved by 5000 iterations of the steepest descent algorithm, systems were equilibrated under NPT conditions. Production runs were performed with a time step of 20 fs updating the neighbor list every 10 steps. For GROMACS we used the V-rescale thermostat^45^ keeping pressure at 1 bar with the Parrinello-Rahman barostat.^46^ In the case of AMBER simulations systems were coupled to a Langevin thermostat^47^ with a collision frequency of 5 ps^−1^ and to a Berendsen barostat^48^ with a relaxation time of 8 ps. In both cases, simulations were run under semi-isotropic conditions. Nevertheless, in our experience simulations are not critically sensitive to the use of particular thermostats, barostats nor moderate variations in their coupling constants.

To characterize structural and dynamical properties, pure DPPC, POPC and POPE patches composed by 64 molecules per leaflet were simulated through 1 μs at four different temperatures (303 K, 313 K, 323 K and 333 K). Each system was simulated by triplicate using different initial random velocities. The area per lipid was obtained by dividing the xy area of the simulation box by the number of phospholipids at one leaflet of the bilayer. The bilayer thickness was measured as the distance between phosphate peaks from density distribution profiles. Second rank acyl order parameters were obtained from the θ angle between the normal of the bilayer surface and the vector corresponding to each bond in the acyl chains using the equation: P_2_ = ½ (3 cos^2^ θ − 1).^10^ Lateral diffusion was computed with *gmx msd* by measuring the mean squared displacement (MSD) of the phosphate beads on the xy plane over the last 0.5 μs of trajectory. Values were averaged among triplicates.

The self assembly capability of the model was tested by randomly putting 140 POPC molecules inside a 6.4 × 6.4 × 11 nm box solvated with 644 WT4 molecules, 20 Na^+^ and 20 Cl^−^ CG ions. The system was simulated for 0.1 μs at 303 K under isotropic NPT conditions and then switched to a semi-isotropic coupling for 1μs.

To study membrane proteins, we generated two classes of bilayer patches formed by 320 lipids per leaflet. One membrane model was made of pure DMPC and the other was asymmetrically composed by an extracellular/cytosolic-like leaflet of POPC and an intracellular/luminal-like leaflet of POPE:POPS at a 65:35 % proportion, roughly representing a typical mammalian cell.^49^ To ensure a well-stabilized membrane support, 1 μs long simulations were performed using the same protocol described above for pure lipid bilayers. These stabilized membrane patches were then employed to embed the following protein systems: i) Staphylococcal α-hemolysin (PDB id: 7AHL); ii) Bacterial outer membrane protein OmpX (PDB id: 2M06); iii) Phospholamban pentamer (PDB id: 2KYV, model #1); and iv) Sarcoendoplasmic reticulum calcium ATPase (PDB id: 5XA7). Proteins were protonated at pH 7 using the pdb2pqr server ^50^. Fine grain structures were converted to CG using SIRAH Tools.^16^ Due to the mushroomlike shape of the Staphylococcal αHL with singular water cavities, extra precaution was needed for its solvation. First, the heptameric structure was solvated in the absence of a phospholipid bilayer and later equilibrated imposing 1000 kJ mol^−1^ positional restraints over the protein backbone of the soluble part and all the beads in the transmembrane region. Finally a POPC/POPE:POPS bilayer was added to the solvated system, removing water molecules that overlapped with the membrane. The orientation of Staphylococcal αHL, OmpX and PLB was set according to the OPM database. After removing the lipids in close contact with the TM domains, two 15 ns equilibration steps were performed. In the first step, positional restraints of 100 kJ mol^−1^ in z coordinate were applied to phosphate head beads, allowing the lipids to occupy the surroundings of the protein without loosing its characteristic thickness. At this stage, proteins were also restrained by 1000 kJ mol^−1^ in x, y and z coordinates. Once the area per lipid of the system was recovered, a second equilibration step was held restraining x, y and z coordinates of lipids’ phosphates and protein backbone ^51^ by 100 kJ mol^−1^, to relax the contacts between protein side chains and lipid tails. Production runs were carried out at 310 K in the absence of any restraint. The complete analyses of the trajectories including tilt angle and density profiles were performed with the tools *gmx gangle* and *gmx density* available in the GROMACS package. Protein-phospholipid salt-bridge formation and occupancy graphs were measured employing VMD analysis tools.^52^ Secondary structure analysis was performed with SIRAH Tools.^16^ Contact conservation and accuracy was measured using the same criteria as in^13^.

## RESULTS AND DISCUSSIONS

### Pure lipid bilayer simulations

It is widely accepted that tilt, distribution and even function of transmembrane proteins are directly affected by the structural properties of the bilayers in which they are embedded.^53^ This is why an accurate reproduction of structural and dynamic features of lipid bilayers becomes crucial when assessing protein-membrane interactions by means of MD simulations. Of paramount importance is the correct and unbiased reproduction of the hydrophobic effect that ultimately dictates the organization of membranes exposing polar heads to the aqueous solvent. Density profiles along the bilayer normal for a DPPC bilayer simulated at 303K suggests that MD simulations using SIRAH implementation result in the correct organization of hydrophilic and hydrophobic moieties (Figure 2A), including the limited water insertion up to the carbonyl region.^54^ Analogous profiles for POPC and POPE are shown in supplementary Figure S1.

**Figure 2.**
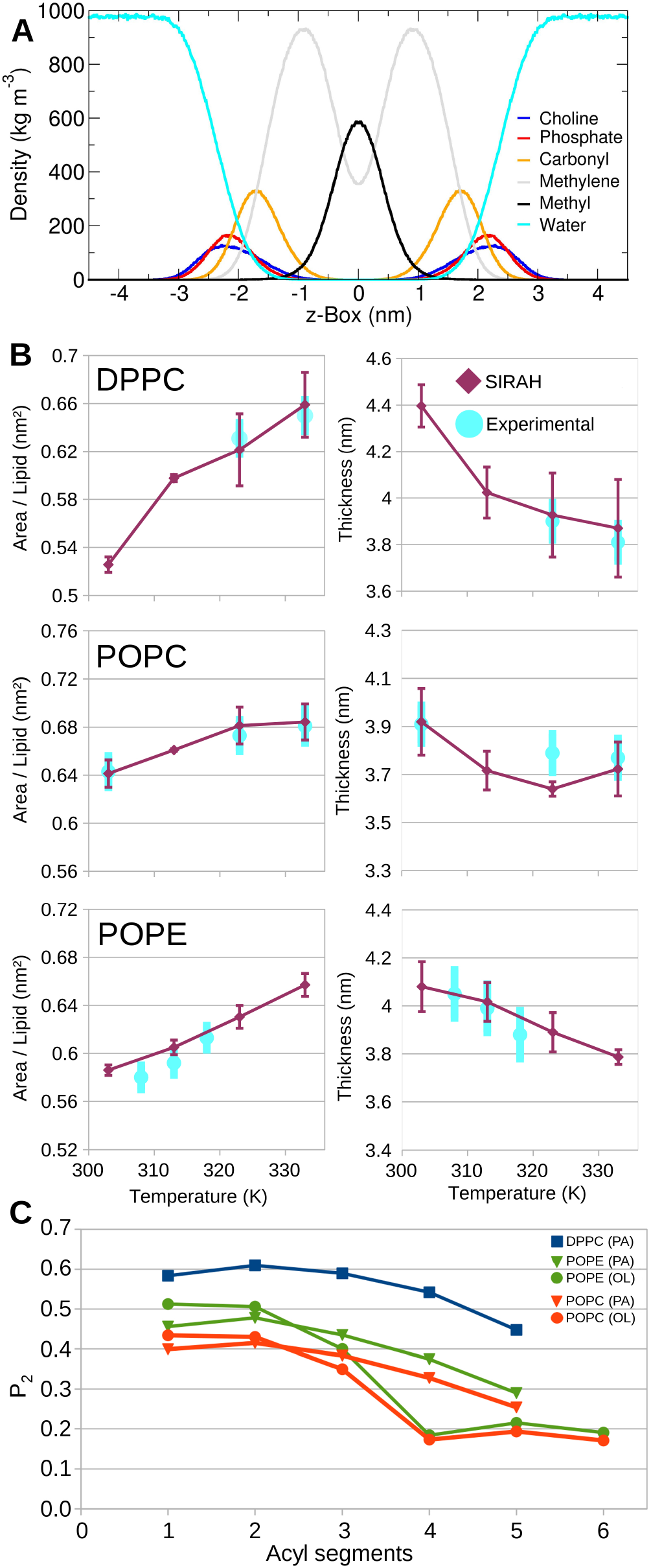
MD simulations of pure lipid membranes. (A) Density profile per moiety of a DPPC bilayer along its normal axis. Analogous profiles for POPC and POPE are shown in supplementary figure S1. (B) Overall thickness and area per lipid are plotted as a function of the temperature for DPPC, POPC and POPE. Despite lacking experimental data at some temperatures, simulations were performed on the same temperature range for completeness. (C) Second rank order parameters per acyl segment at 303 K. Simulations were performed in GROMACS.

The overall thickness of the simulated DPPC, POPC and POPE membranes can be estimated from the distance between the peaks of the phosphate distribution in Figures 2A and S1. These measurements are directly comparable to experimental results from Small-Angle Neutron and X-ray Scattering obtained by Kučerka et al. at different temperatures.^29,56^ As Figures 2B shows, a correct description of the membrane thickness as a function of temperature is observed for the three species. In particular, the decrease in membrane thickness when raising the temperature from 303 K to 333 K is well reproduced within the reported experimental error. As expected, the progressive reduction in the thickness is accompanied by an increase in the area per lipid in the same range of temperatures (Figure 2B). In the case of the SIRAH implementation for AMBER we also observed comparable values and trend in the temperature-dependent thickness and area per lipid values (supplementary Figure S2).

The effect of temperature over the lipid structure could be also followed at a higher detail by measuring order parameters in the esterified acyl tails. To this end, the second-rank order parameter was computed for consecutive CG methylene beads. Perfect alignment, perfect anti-alignment and a random orientation with the bilayer normal is indicated by P_2_ equals 1, −0.5 and 0 respectively. It is important to remark that, as a consequence of the coarsening, these values can only be tested against other CG models,^57^ being just qualitatively comparable with experimental data. As seen from Table 1, the average tail order parameter follows the expected decrease with the temperature. A more detailed inspection indicated that P_2_ values gradually decreased for saturated acyl tails when moving towards the terminal beads. DPPC bilayers displayed the highest order, as it could be expected for saturated species. The lowest values (0.32 ± 0.01) were found for POPC at 303K, showing good agreement with the average tail order value of 0.34 ± 0.01 reported in the popular MARTINI force field (Table 1).^10^ Despite having the same types of acyl tails, POPE showed higher P_2_ values. This can be explained by the smaller size of ethanolamine compared with choline head groups, allowing a closest packing as evidenced from lower area per lipid values^58^ (Figure 2B). Moreover, our lipids described correctly the expected decrease in order along the lipid tail with a shoulder at the oleoyl unsaturation (Figure 2C).

**Table 1.** Average tail order parameter (P_2_)

|  | 303 K | 313 K | 323 K | 333 K |
| --- | --- | --- | --- | --- |
| DPPC | $0.55 \pm 0.01$ | $0.43 \pm 0.01$ | $0.40 \pm 0.03$ | $0.36 \pm 0.02$ |
| POPC | $0.32 \pm 0.01$ | $0.30 \pm 0.01$ | $0.28 \pm 0.01$ | $0.26 \pm 0.02$ |
| POPE | $0.37 \pm 0.01$ | $0.34 \pm 0.01$ | $0.30 \pm 0.01$ | $0.29 \pm 0.01$ |

Lipid lateral diffusion coefficients (D_L_) are broadly employed to estimate the dynamics and fluidity of biological membranes. One of the most widely used techniques to measure D_L_ is Fluorescence Recovery After Photobleaching (FRAP) that can calculate self-diffusion of lipids within time windows of few hundreds of nanoseconds. This time scales are compatible with MD simulations making this experiments a good control to assess the capacity to reproduce the membrane dynamics^59^. Experimental D_L_ values from 13 μm^2^ s^−1^ at 318 K^60^ to 8 μm^2^ s^−1^ at 323 K^61^ have been reported for DPPC. Lateral diffusion coefficients for POPC collected from the literature by Pluhakova et al.^18^ at shows how diffusion coefficients vary between 4 to 5 μm^2^ s^−1^ and 7 to 16 μm^2^ s^−1^ at 300 K and 320 K, respectively. For the case of POPE there is scarce information, finding only one value of 5.5 μm^2^ s^−1^ at 308 K.^62^ In the present work, D_L_ was computed using the Einstein relation based on the average lateral mean squared displacement (MSD) which was calculated on the position of the phosphate beads. As a first observation, the SIRAH lipids diffuse faster than atomistic ones. This is a common effect observed in CG approaches, derived from the lumping of degrees of freedom, which translates in smoother free energy landscapes, and reflects in faster diffusion rates.^63^ In our case, using a scaling factor of 4 (the same reported for MARTINI^57^) resulted in a good comparison between experiments and simulations (Tabla 1). Indeed, the correct temperature dependence was recovered.

**Table 2.** Lateral diffusion coefficient (μm^2^ s^−1^)^a^

|  | 303 K | 313 K | 323 K | 333 K |
| --- | --- | --- | --- | --- |
| DPPC | $6.1 \pm 1.5$ | $8.4 \pm 0.8$ | $11.2 \pm 1.3$ | $14.2 \pm 2.7$ |
| POPC | $5.7 \pm 0.5$ | $8.3 \pm 0.8$ | $12.2 \pm 1.2$ | $13.9 \pm 2.6$ |
| POPE | $2.6 \pm 0.6$ | $5.1 \pm 0.9$ | $7.2 \pm 1.4$ | $8.8 \pm 0.4$ |

<sup>a</sup> Shown values are divided by a factor of 4.

### Self assembly simulations

To complete the validation of our CG parameters for lipids, we assessed their spontaneous self-assembly capacity. To this aim, POPC molecules were randomly placed in a rectangular box and then solvated with WT4 and 150 mM of Na^+^ and Cl^−^. After 0.05 μs of simulation at. 303 K under isotropic conditions, polar heads started to group and hydrophobic tails established the first interactions through the periodic boundary conditions forming a transient water pore that disappeared at about 0.1 μs (Figure 3A-C). After that time a flat bilayer was completely formed (Figure 3D). To control the structural properties of the spontaneously formed bilayer, the simulation was extended for other 0.9 μs using a semi-isotropic pressure coupling. Area per lipid and thickness were measured and averaged over the last 0.1 μs of simulation resulting in 0.62. ± 0.02 nm^2^ and 3.9 ± 0.1 nm, respectively, in close proximity with experimental values^55^ as well as previous simulations started from well constituted bilayers (Figure 2B).

**Figure 3.**
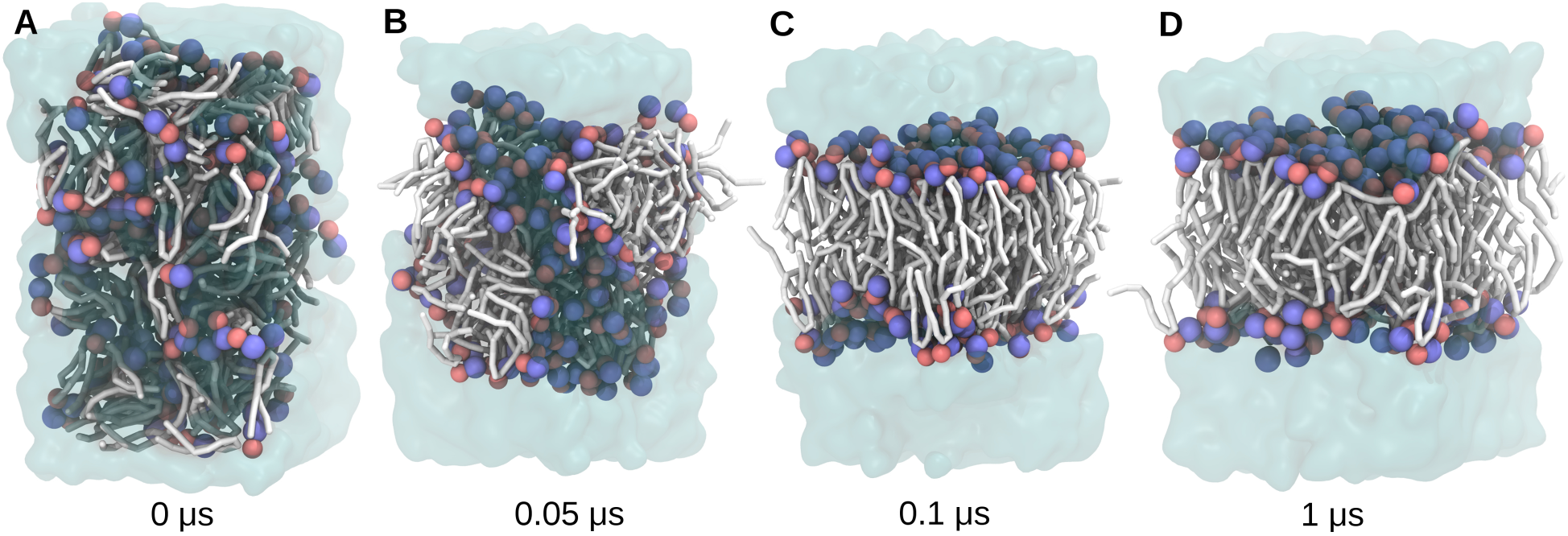
Representative snapshots of the POPC self-assembly simulation. Choline and phosphate beads are represented in blue and red spheres, respectively, while the rest of the molecule is drawn in white tubes. Water molecules are shown as a transparent-cyan surface. Panels A to D indicate different time points along the simulation.

### Membrane proteins

Besides achieving a correct reproduction of lipid bilayer properties, the main goal of the present contribution is to provide a good framework for the simulation of membrane proteins. To this aim we choose four particularly challenging systems as study cases, highlighting the potentialities and limitations of our approach. A series of structural properties were analyzed in order to validate the correct selection of parameters, particularly those involved in the protein-lipid interactions. The analysis that will be described in detail in this section also included secondary structure reproduction, contact conservation, RMSD values and transmembrane (TM) domain tilting. The availability of experimental data about orientation in membrane and lipid protein-interactions for the last two systems addressed (SERCA and PLB) enabled us to perform more detailed comparisons with simulation results.

### Case study 1: Staphylococcal a-hemolysin

The bacterial protein α-hemolysin belongs to the family of pore forming toxins. These are soluble proteins that upon oligomerization form transmembrane pores that permeate ions with poor specificity generating cellular osmotic imbalance and cell lysis.^64,65^ The structure of this heptameric complex obtained by X-ray experiments^66^ presents a mushroom shape with an extracellular cap domain composed by 7 β-sandwiches, a stem domain composed by 14 antiparallel β-strands, and seven rim domains partially inserted into the membrane. CG simulation were preformed in a plasmatic-like membrane of POPC/POPE:POPS. A special solvation protocol (see Methods) was employed to fill the water cavities located at the central pore and the rim domain. This precaution has been previously considered by Aksimentiev and Schulten in their fully atomistic simulations of αHL in a DPPC bilayer.^67^

For structural stability determination, RMSD was calculated for each monomer and averaged over the last 0.1 μs giving values of 0.45 ± 0.02 nm. Secondary structure was monitored on the TM region. The initial β-strand content in the X-ray structure descended from 91 to 76 ± 2 % after 2 μs of unrestrained simulations (Figure 4A). This partial structural loss was observed at the trans end of the β-barrel, near the solvent-exposed loops facing the intracellular lumen (see insets in Figure 4A). The wide internal diameter of the pore in α-HL permitted its solvation by CG water, which contributed to maintain the barrel shape of the stem domain. The initial diameter of 2.6 nm, measured from Cα atom in the crystallographic structure, was conserved with small deformations towards an oval shape observed after 1.3 μs that was partly recovered in the last 0.2 μs of simulation (Figure 4B). All these features were present in the results reported by Aksimentiev and Schulten^67^ using timescales of 0.1 μs. RMSD values of 0.28 nm, structural distortions in the trans end of the β-barrel, and the adoption of an ellipsoidal shape of the pore are quantitatively well compatible with our values, shown in Figure 4B for the CG simulation. Similar shape deformations of α-HL were also observed in more recent atomistic ^68^ and CG simulation^69^ within POPC and DMPC membranes, respectively. In both cases a pronounced collapse at the trans face of the pore is generated according to the lipid composition of the membrane.

**Figure 4.**
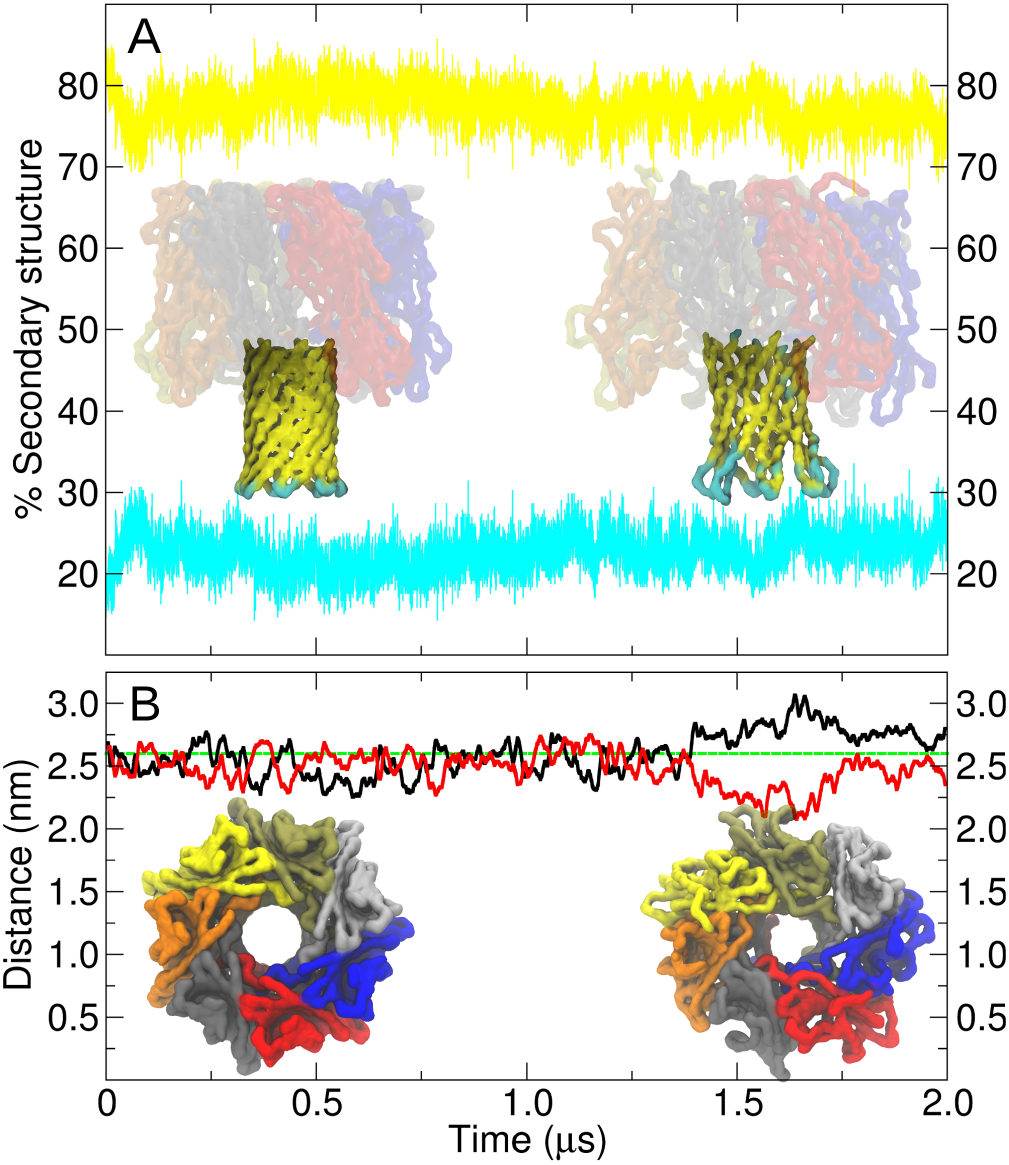
Simulation of α-hemolysin in a POPC/POPE:POPS bilayer. (A) Secondary structure content as a function of time for the stem region (β-sheets in yellow, coils in cyan). Insets show side views of initial (left) and final (right) structures. Molecular drawings represent the backbone of the protein. The stem region is colored by secondary structure, while extracellular domains are colored by chain. (B) Pore diameter along the 2 μs trajectory. The value obtained from the X-ray structure is colored in green, major and minor diameters observed during the simulations are represented by black and red traces, respectively. Extracellular views of the initial and final structures are colored by monomers.

### Case study 2: Bacterial outer membrane protein OmpX

In the previous paragraph we showed a good reproduction of structural features in a relatively large proteinaceous pore. Similarly good results had also been obtained for SIRAH in the simulation of the pore formed by the hexameric Connexin 26 hemichannel.^70^ However, in both cases, these multimeric complexes feature large channels with diameters over ~1.2 nm, which are experimentally known to conduct solvated ions and small molecules. Such molecular topologies are well suited for CG approaches as bulky CG water can permeate through the pore solvating them completely. As discussed for α-HL, a proper solvation is crucial for a good reproduction of the structural and dynamical properties of proteins. Therefore, we sought to stress test our force field in a situation in which the protein features a smaller orifice that precludes WT4 water to penetrate. With this aim we selected the bacterial outer membrane protein OmpX. This β-barrel protein belongs to a family of highly conserved proteins that exerts virulence by neutralizing host defense mechanisms.^71^ Its structure, resolved by solution NMR techniques,^72^ consists of 8 antiparallel amphipathic β-strands with hydrophilic surface-exposed loops and periplasmic turns (Figure 5A and B).^73^ Alike α-HL, OmpX was embedded in a mixed lipid patch composed by POPC and POPE:POPS leaflets. The average RMSD from the last 0.1 μs of simulation was 0.60 ± 0.03 nm, while native contacts were conserved in 72 ± 1% with 81 ± 1% accuracy. The β-sheet content dropped from 76% to 57 ± 3% during the simulation. These conformational changes were likely the result of a constriction at the interior of the barrel (Figure 5C and D). These cavities would normally be filled with water but due to the coarse nature of the model, the interior of the protein could not be properly solvated leading to the lost of the characteristic twist in their strands. This example is presented as a limitation that should be considered when choosing CG approaches to simulate TM proteins with narrow water-filled cavities or other features, which are crucially depending on fine or atomistic details.

**Figure 5.**
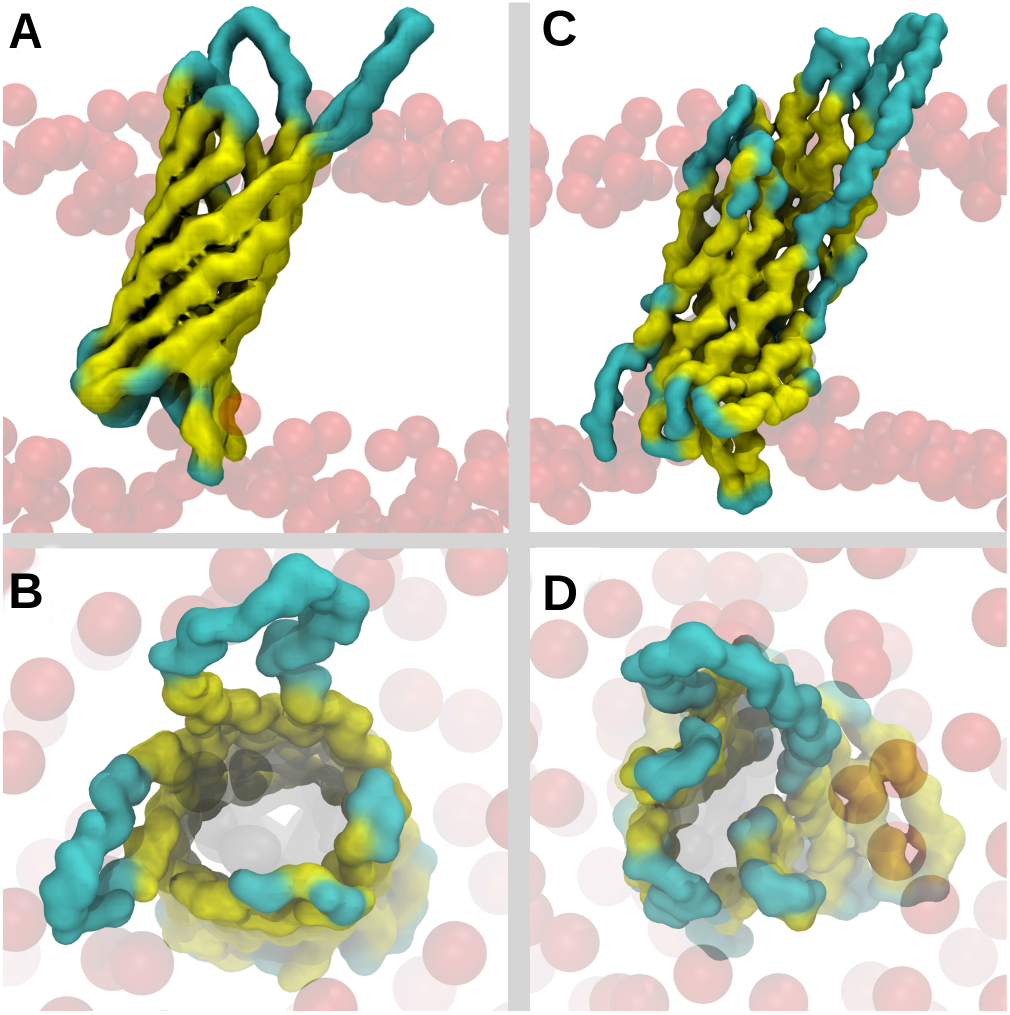
(A) Initial structure of the OmpX protein embedded in a POPC/POPE:POPS bilayer. Backbone is colored by secondary structure (cyan = coil; yellow = β-sheet). Phosphate beads are shown with red transparent van der Waals spheres. (B) Cytosolic view of the initial structure of the β-barrel. (C,D) Same as panels A and B for the final conformation after 2μs.

### Case study 3: Phospholamban pentamer

Phospholamban (PLB) is a type II membrane protein that inhibits the SERCA pump, regulating calcium homeostasis in cardiac muscle. PLB is composed by 52 amino acids divided into domain I (cytosolic residues 1 to 30) and domain II (TM residues 31 to 52). Both domains feature a predominant α-helix conformation. Experimental information regarding protein orientation is available for both PLB and SERCA, providing the opportunity to check the sensibility of our force field to different characteristics of the membrane, in particular its thickness. Moreover, the pentameric arrangement of PLB offers a good test case to evaluate protein-protein interactions and stability within lipid bilayers. The pentameric structure of PLB, as obtained by a hybrid in-solution and solid-state NMR method^74^ (PDB id: 2KYV), is characterized by a “pinwheel” conformation stabilized by the adsorption of domain I from each monomer to the cytosolic leaflet surface. Such conformation was well maintained along a 2μs trajectory in a POPC/POPE:POPS membrane, as it could be inferred from the density profile and the trajectory’s final snapshot shown in Figure 6A and B. The stability of the pentamer was estimated from RMSD values obtained over the last 0.1 μs when compared with model #1 of the NMR family 2KYV. Analysis was performed separately for both domains. The tight packing within the membrane environment resulted in a rather stiff conformation of domain II with RMSD of 0.23 ± 0.12 nm. Additionally, average native contact conservation and accuracy over the last 0.1 μs of simulation were 91 ± 3 % and 93 ± 5 %, respectively. Both values were indicative of high stability in the reciprocal conformations of the five independent helices. On the other hand, the solvent exposed domain I reached higher RMSD values of 1.26 ± 0.65 nm. Nonetheless, calculation over individual segments resulted in considerably lower values for domain I (bellow ~0.5 nm, see supplementary Figure S3), being the large structural fluctuation partly explained by the adaptation of the helices to the undulation of membrane. This behavior is also observed in the family of structures obtained by NMR reaching RMSD values up to ~ 0.9 nm.

**Figure 6.**
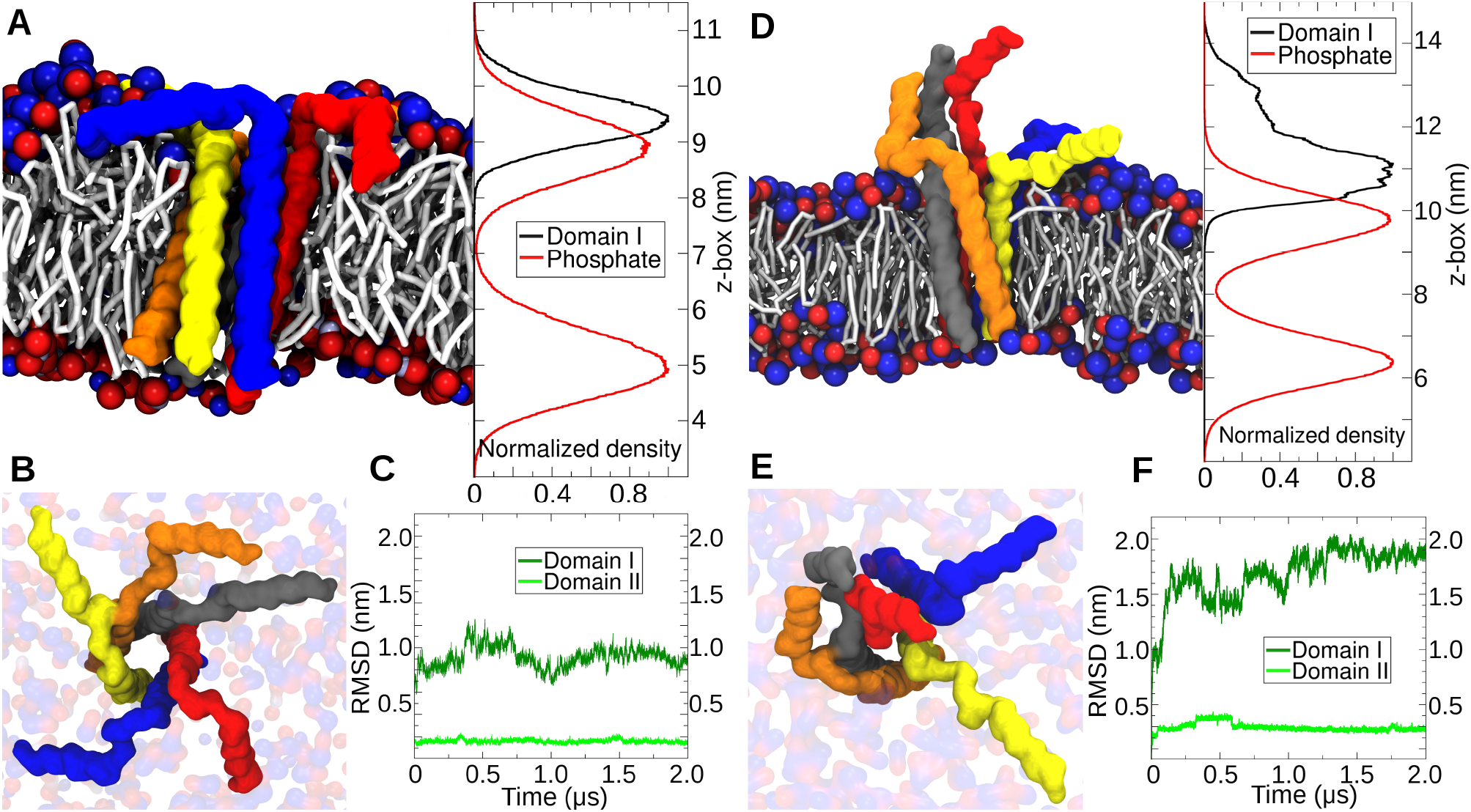
(A) Final snapshot from a 2 μs trajectory and normalized density profile across the z-box for the cytosolic helices of PLN pentamer and phosphate beads in a POPC/POPE:POPS bilayer. PLN backbone is colored by chain, while lipid head groups are colored by charge (blue: positive, red: negative). (B) Top view of the final snapshot in panel A. (C) Time evolution of RMSD values for the five cytosolic (domain I) or TM (domain II) regions of the PLN pentamer, taking as reference the model #1 from NMR structure 2KYV. Values for single replicas are shown (D-F) Same as panels A-C for a DMPC bilayer.

The tilt of TM helices reduces the hydrophobic mismatch between the protein and the lipid bilayer, contributing for their stability.^75^ For PLB pentamer, this property was studied by all atom MD simulations using DOPC^74^ and DOPC:DOPE bilayers^76^ showing tilt angles in the range of 5° to 25° with maximum at 11° and 15° of each distribution, respectively. Taking into account the higher sampling time explored here, our results were in very good agreement with the atomistic MD. Measuring the angle between the vector formed by Leu 28 and Ile48 Cα beads respect to the normal of the simulation box showed that PLB tilted preferentially around 17° with a continuous dispersion of values from 5° to 35° (supplementary Figure S4). Another important quality index for a structurally unbiased force field, as SIRAH, is the conservation of secondary structure elements of integral membrane proteins.^63^ Capping α-helices of soluble proteins and peptides lack intra-helical hydrogen bonds stabilizing the first/last four amine/carboxylic groups of the backbone.^77^ However, in TM proteins, helix ends can find Hydrogen bonding partners within the polar head groups of the bilayer.^53^ Here, we observed that the phospholipidic environment completely stabilized C-terminal TM helices along the 2 μs trajectory, while solvent exposed N-temini were prone to lose their conformation (supplementary Figure S5).

Next, we sought to study the sensibility of the force field to changes in the membrane properties. We embedded the PLB pentamer in a DMPC bilayer, which presents a thickness of ~ 3.3 nm at 310 K.^78^ In line with our expectations, phospholipid composition impacted in the behavior of the cytosolic helices and sporadic detachment from the bilayer surface was observed during the trajectory (Figures 6D, E). This conformational behavior has been associated to a “bellflower” topology^79,80^; or in the case of PLB monomers to the “R” state.^81^ In the R state, domain I of PLB is more dynamic, solvent exposed and prone to adopt unstructured conformations. Worthy, this has also been found with Magic-Angle-Spinning solid state NMR determinations employing DMPC bilayers.^82^ This conformational flexibility has been directly correlated with its regulatory function upon interactions with the SERCA pump.^83,84^ Despite the dynamical character of domain I in DMPC (RMSD = 1.80 ± 0.15 nm), structural stability of domain II was conserved, displaying RMSD values of 0.36 ± 0.08 nm and conservation and accuracy of native contacts of 86 ± 3 % and 89 ± 2 %, respectively. For both POPC/POPE:POPS and DMPC systems, RMSD and contact values were averaged and its standard deviations calculated over the last 0.1 μs of three independent replicas.

To further test the correct reproduction of protein-protein interactions in phospholipidic environments we introduced Cys36Ala, Cys41Phe and Cys46Ala mutations in the pentameric structure. This sequence modification, also known as PLB^AFA^, favors the monomeric state of Phospholamban increasing its capacity to inhibit SERCA.^85^ We embedded the mutant pentamer in a DMPC bilayer and observed, early in the equilibration steps, how the bulkier Phe41 protruded towards the lipid region unlike the wild type Cys41 that locates in the hydrophobic protein interface.

After ~ 0.1 μs a DMPC molecule introduced its acyl chain in the small cavity generated by Cys41Phe (Supplementary movie M1) starting a destabilization process that resulted in an almost complete dissociation of one monomer, except for its C-terminal domain. The triple mutation generated major structural perturbations over domains I and II reflected in RMSD values of 2.23 ± 0.28 nm and 0.57 ± 0.03 nm, respectively (Supplementary figure S6). This highlights the sensibility of the force field to point mutations.

### Case study 4: Sarcoendoplasmic reticulum Calcium ATPase (SERCA)

The SERCA pump has been extensively studied and crystallized along several states of its catalytic cycle. It was the first membrane protein in which the head groups of the phospholipid bilayer were clearly visible from the electron density,^86^ leading to the measurement of the tilt angle between the protein and the membrane. Electron density maps of four SERCA intermediate states showed a thickness-driven tilting. Specific interactions between phospholipids and Lysine, Arginine or Tryptophan residues were also recognized to participate in the orientation of the protein in the membrane, which also have functional roles in the dynamics of the pump. An additional challenge is the presence of two bound Calcium ions at the TM region of the enzyme, which structural role is still matter of debate at experimental and computational level.^87^

To assess the influence of bilayer’s thickness in the tilt of SERCA, the E1-2Ca^++^ state (PDB id: 5XA7) was embedded in POPC/POPE:POPS and pure DMPC bilayers. Aimed to challenge the capability of SIRAH to sample the conformational space, TM helices were initially placed perpendicular to both bilayer patches and the tilt angle between a vector formed by Cα of residues Val790 and Thr805 (TM helix #5) and the normal of the simulation box was followed through 3 μs (Figure 7A). In both cases the protein spontaneously changed its orientation, but when it was embedded in the plasmatic-like membrane it experienced a sensibly less marked change, maintaining its orientation at values lower than 30°, with maximum probabilities between 17° and 26° (Figure 7B). In contrast, the same protein embedded in the DMPC bilayer rapidly tilted to values up to 55° and after 1 μs of simulation stabilized around 35°, i.e., matching the experimental value determined from X-ray crystallography (Figure 7B, black trace). Non of these protein movements affected the thickness of the membrane patches. Indeed, averaging thickness values over the las 0.1 μs of simulation resulted in 3.9 ± 0.1 nm and 3.3 ± 0.1 nm (Figure 7C), respectively.

**Figure 7.**
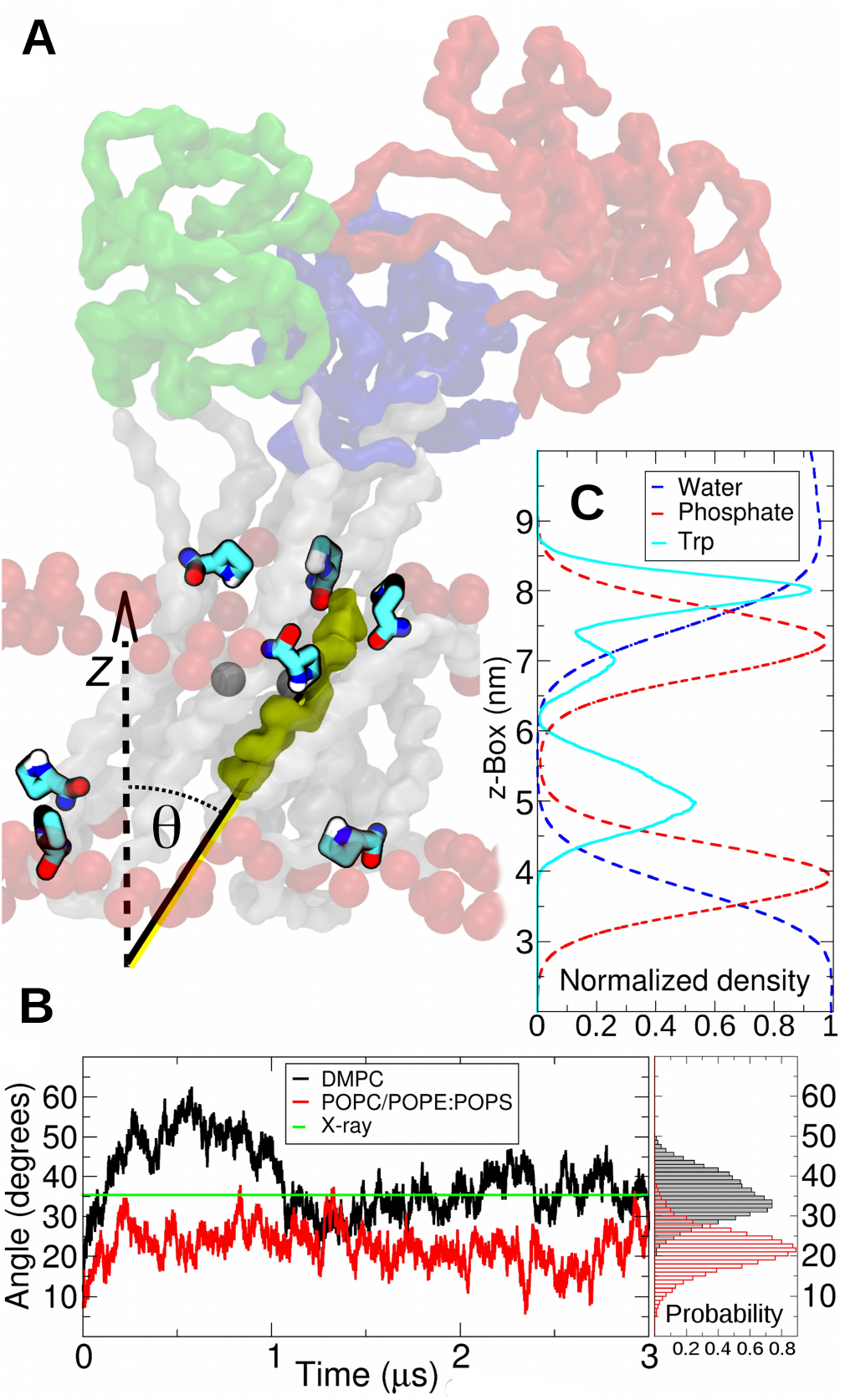
(A) Representative snapshot of the SERCA E1-2Ca++ state embedded in a DMPC bilayer. Tryptophan residues are represented by tubes and colored by element and phosphate beads by red spheres. The TM helix 5, used to measure the tilting angle of the protein is highlighted in yellow. Calcium ions are shown in dark gray as a reference. (B) Comparison of the X-ray determined tilt angle of the TM domain of SERCA and those obtained with SIRAH for two membrane compositions. Probabilities are calculated from the last 2 μs of simulation. (C) Normalized density profile showing the distribution of Tryptophan side chains (cyan), phosphate beads (red) and water (blue) across the z-box.

Subsequently, we wondered if the tilt was sensitive to the initial conditions. For this, we started a second simulation orienting the SERCA in a DMPC bilayer according to the electronic density data from X-ray experiments. DMPC was chosen because of its thickness similarity to the 3.1 nm bilayer employed in the X-ray crystallography.^86^ Again, the crystallographic tilt was nicely reproduced within a 5 μs simulation time (Supplementary Figure S7).

In sight of the good agreement with the experimental data, and despite the coarseness of our approach, we attempted to characterize the molecular determinants causing the conformational behavior of the protein. This longest simulation facilitated identifying a series of Tryptophan residues stabilizing interactions between the protein and the membrane surface. It is known that Tryptophan residues at the interface of integral membrane proteins behave as “floaters” with their polar side facing to the water and their hydrophobic regions dipped into the bilayer.^88^ In agreement with these observations, we found that Trp50, 77, 107, 288, 832, 855, and 932 formed a belt that modulate the orientation of the TM domain (Figure 7A). These amino acids were originally identified from the X-ray data and their interaction was fully maintained during the simulation, as reflected by their sustained occupancy in that region (Figure 7C).

In addition to Tryptophan floating, two different types of interactions between phospholipids and single amino acids have been proposed^86^: i) Snorkeling, when basic side chains emerge from the hydrophobic region of the bilayer towards the solvent forming stable salt-bridges with phosphate groups; and ii) Anchoring, in this case basic residues located at the water phase orient their side chains towards the membrane surface, interacting with a higher number of phospholipids. To analyze these interactions, we traced back lipid molecules able to transiently form salt-bridge interactions with Lysine or Arginine residues at the protein-membrane interface during the 5 μs trajectory in DMPC. To study snorkeling, we focused on Lys262, located at the hydrophobic region of the TM helix #3 (Figure 8A). During the simulation, this residue preferentially oriented its basic moiety to the cytosolic surface of the bilayer. This was evidenced by the pseudo χ angle of Lys262, which presented a preferential distribution between 120° and 180° (Figure 8B). After 0.6 μs of simulation Lys262 formed a stable salt-bridge with a DMPC molecule for approximately 2 μs (Figure 8C). A second DMPC molecule that formed a salt bridge interaction until the end of the trajectory later occupied this binding site. On the other hand, anchoring events were also verified between basic residues and phospholipids. In particular Arg110, located at TM helix #2 interacted with a series of different polar heads (Figure 8D). In contrast with Lys262 the pseudo χ angle distribution of the side chain for this residue showed three preferential orientations (Figure 8E). In addition to conformations at 160° to 240° pointing to the solvent phase, two other conformations from 80° to 140° and 280° to 320° (i.e., pointing to the membrane), were observed. This multiplicity of conformations was characterized by a constant turn over of phospholipids forming salt-bridges through the simulation (Figure 8F).

**Figure 8.**
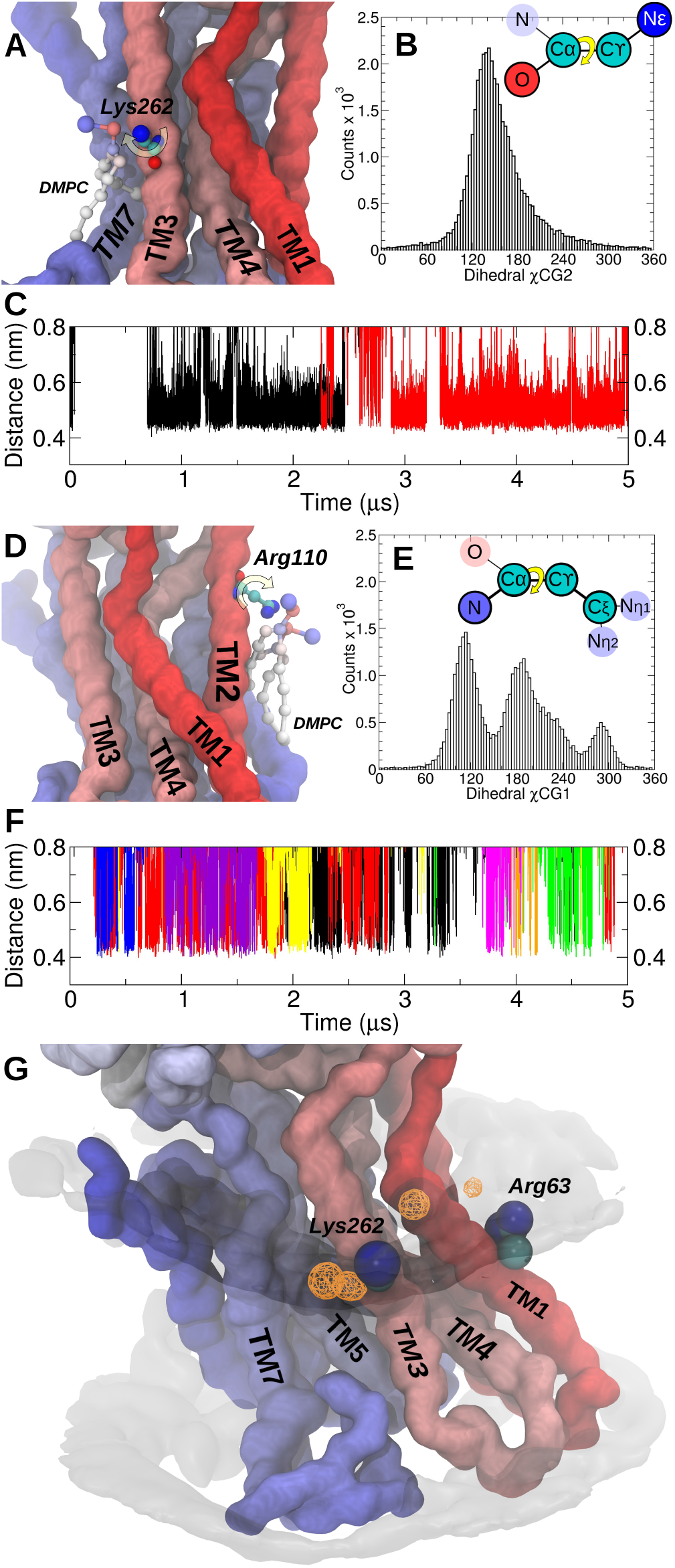
(A) Molecular representation of the snorkeling interaction between Lys262 and a DMPC molecule. Interacting moieties are represented by balls and sticks. The protein backbone is represented with surfaces and colored using a RWB color scale to distinguish each TM helix. (B) Distribution of pseudo χ angle of Lys262. The inset shows a schematics of the measured dihedral at CG level. Beads are named according to the mapped atom. (C) Tracking of salt-bridges between Lys262 and phospholipids. Time evolution of distance between the basic moiety of Lys262 and any phosphate bead able to approach closer than 0.5 nm during the simulation. Individual interacting molecules correspond to different color traces. (D) Anchoring of Arg110 to two DMPC molecules using same representation from panel A. (E, F) Same as panels B, C for Arg110. (G) Occupancy analysis indicating the annular shell (gray transparent surface, isovalue = 10%) and snorkeling lipid-protein interactions (orange mesh, isovalue = 50%)

The analysis of the trajectories suggest that this dissimilar behavior between Lys262 and Arg110 could be related to specific lipid organization around TM proteins. Through an occupancy analysis of the phosphate groups, annular and non-annular binding lipids were observed during the 5 μs simulation (Figure 8G). Setting an occupancy threshold of 10% resulted in a ring-shaped surface around the TM domain while increasing this value to 50% displayed very circumscribed regions in the surroundings of Lys262 and Arg63 basic moieties, both previously reported as implicated residues in snorkeling interactions.^86^ For the case of Arg110, interacting DMPC molecules correspond to the annular lipidic shell, a region characterized by surrounding the TM domains for periods of time of 0.1-0.2 μs having little structural specificity.^89^ Alternatively, the DMPC molecule bound to Lys262 would correspond to a non-annular site, known for being located between TM α-helices tightly binding lipid molecules.^75^ In our simulation protocol this corresponded to the hydrophobic pocket formed by TM helices #3, #5 and #7. It is interesting to notice that this region is very well characterized as the binding site of thapsigargin, a potent inhibitor of SERCA.^90^

## CONCLUSIONS

A new set of CG phospholipid models compatible with the SIRAH force field is presented. Along with native porting to the popular MD engines GROMACS and AMBER, which are freely available from the web site www.sirahff.com. To the best of our knowledge, this is the only available Coarse-Grained force-field ported to AMBER and able to handle simulations containing DNA, explicit solvent, ions, proteins and membrane as provided in the present SIRAH package. The use of long-range electrostatics and structurally unbiased protein description is another distinctive feature that enabled the description of electric field driven conformational transitions in membrane proteins.^70^ The current implementation includes both residue-based and fragment-based topologies describing residues DMPC, DPPC, POPC, POPE and POPS, head groups PC, PE and PS, and tails MY, PA and OL. Although we presented here only five molecular species, they provide a minimalist representation of the phospholipids variety including short/long and, saturated/unsaturated tails, as well as polar and acidic heads. Moreover, the use of fragment-based topologies offers the chance to considerably expand the combinatorial space actually available in the residue-base library of lipids arriving to 27 different species. In addition, the proposed mapping scheme provides a simple recipe for new lipid tails. In that sense, special attention was devoted to the user-friendliness of SIRAH by providing compatible MAP files to most used all-atoms and united-atoms force fields. Following the SIRAH’s strategy, parameters were assigned to reproduce experimental structural data such as area per lipid, overall thickness and average tail order. These properties showed quantitatively correct sensitivity to thermal variation in pure lipid bilayers. The correct hydrophobic/hydrophilic balance was assessed by the self-assembly capacity of the model. It is worth to notice that the absence of topological restraints in the protein models may demand extra precaution in the assignment and control of lipid-protein interactions. To avoid overestimating the hydrophobic attraction between terminal acyl beads and proteins’ beads, these interactions were calculated outside the combination rules. In a similar way, salt-bridge formation between lipid heads and charged amino acids was set following the same criteria used for proteins.^13^ Simulated TM proteins correctly reproduced both α-helical and β-sheet secondary structure elements. RMSD values showed a well structural reproducibility, being in the range of water-soluble peptides previously reported in the last version of the force field. Proteins showed the capacity to spontaneously adapt their conformations to reduce the hydrophobic mismatch in bilayers with different overall thickness. As discussed, the behavior of integral membrane proteins is not only guided by roughly defined polar/apolar environments but also by specific interactions such as Tryptophan floating and Lysine and/or Arginine snorkeling and anchoring, in very good agreement with experimental reports present in the literature. Nevertheless, potential users of SIRAH are encouraged to consider the limitations of the force-field such as the poor structural reproduction of transmembrane pores with small water cavities. The implementation of this new set of parameters and topologies, combined with its predecessor protein and water models, allows for a speed up over two orders of magnitude, profiting from GPU acceleration and without resigning the valuable physico-chemical insight of MD simulations.

## ACKNOWLEDGMENTS

This work was partially funded by FOCEM (MERCOSUR Structural Convergence Fund), COF 03/11. M.R.M. and S.P. belong to the SNI program of ANII. E.E.B. is beneficiary of a postdoctoral fellowship of CONICET (Consejo Nacional de Investigaciones Científicas y Técnicas, Argentina). Some of the graphic cards used in this research were donated by the NVIDIA Corporation.

## SUPPLEMENTARY MATERIAL

*Data not shown*

